# What is a supercoiling-sensitive gene? Insights from topoisomerase I inhibition in the Gram-negative bacterium *Dickeya dadantii*

**DOI:** 10.1101/2021.04.09.439150

**Authors:** Maïwenn Pineau, B. Shiny Martis, Raphaël Forquet, Jessica Baude, Camille Villard, Lucie Grand, Florence Popowycz, Laurent Soulère, Florence Hommais, William Nasser, Sylvie Reverchon, Sam Meyer

**Affiliations:** Université de Lyon, INSA Lyon, Université Claude Bernard Lyon 1, CNRS UMR5240, Laboratoire de Microbiologie, Adaptation et Pathogénie, 69621 Villeurbanne, France; Université de Lyon, INSA Lyon, Université Claude Bernard Lyon 1, CPE Lyon, CNRS UMR 5246, Institut de Chimie et Biochimie Moléculaires et Supramoléculaires, 69622 Villeurbanne, France

## Abstract

DNA supercoiling is an essential mechanism of bacterial chromosome compaction, whose level is mainly regulated by topoisomerase I and DNA gyrase. Inhibiting either of these enzymes with antibiotics leads to global supercoiling modifications and subsequent changes in global gene expression. In previous studies, genes responding to DNA relaxation induced by gyrase inhibition were categorised as “supercoiling-sensitive”. Here, we studied the opposite variation of DNA supercoiling in the phytopathogen *Dickeya dadantii* using the non-marketed antibiotic seconeolitsine. We showed that the drug is active against topoisomerase I from this species, and analysed the first transcriptomic response of a Gram-negative bacterium to topoisomerase I inhibition. We find that the responding genes essentially differ from those observed after DNA relaxation, and further depend on the growth phase. We characterised these genes at the functional level, and also detected distinct patterns in terms of expression level, spatial and orientational organisation along the chromosome. Altogether, these results highlight that the supercoiling-sensitivity is a complex feature, which depends on the action of specific topoisomerases, on the physiological conditions, and on their genomic context. Based on previous *in vitro* expression data of several promoters, we propose a qualitative model of SC-dependent regulation that accounts for many of the contrasting transcriptomic features observed after gyrase or topoisomerase I inhibition.

## Introduction

DNA supercoiling (SC) is the product of torsional stress ubiquitously experienced by the double-helix in all kingdoms of life. In bacteria, the chromosome is maintained in a steady-state level of negative SC by the interplay of nucleoid associated proteins (NAPs) and the activity of topoisomerases. The DNA gyrase (belonging to class II topoisomerases) introduces negative supercoils by ATP-dependent double-strand cleavage; conversely, topoisomerase I (topoI, class IA) removes excessive negative supercoils through ATP-independent single-strand cleavage, and topoisomerase IV (topoIV, class II) through ATP-dependent double-strand cleavage (1–3). Cells react to SC imbalances by a fast adjustment of topoisomerase activities, according to a homeostasis mechanism (4). This balance plays a key role in many cellular functions, and in particular in the expression of the genome, which is our focus in this study.

The presence of torsional stress in the DNA template is known to affect the transcription process at several successive steps: by modulating the binding of transcriptional regulators and RNA Polymerase (RNAP) itself, the formation and stability of the open complex (5), promoter clearance (6), elongation and termination (7, 8). As a result, SC was soon recognized to act as a global transcriptional regulator (9, 10), although the precise underlying mechanisms remain controversial and subject to many complicating parameters. Early studies demonstrated a strong regulatory action of SC on the promoters of stable RNAs in *Salmonella enterica* and *Escherichia coli* (5, 11), pointing to a role in growth control (10) consistent with the close relationship between SC and the cell metabolism (12). But other promoters were found to be equally affected (7, 13), which was then confirmed and broadened by high-throughput transcriptomics methods (14–16). In analogy to the “regulons” of transcriptional factors, these promoters were often termed “supercoiling-sensitive”, although that notion remains poorly defined, considering the lack of clearly identified sequence determinants (17), and the variability in the response of many promoters to SC alterations depending on their context and the experimental protocol of the assay. For example, the *lacP* promoter of *E. coli* is strongly repressed by DNA relaxation *in vitro* (7), but is unaffected *in vivo* (14, 15); the proportion of genes activated by DNA relaxation in *S. enterica* varied between 70% in a random fusion assay (18) and 27% in a RNA-Seq transcriptome (19).

*In vivo*, these responses to SC variations were obtained by two distinct methods (20). The expression level can be measured in topoisomerase mutant strains, which usually exhibit a different SC level than the parental strain cultured in the same conditions (20, 21); however, the difference in promoters’ expression then reflects not only the direct regulatory effect of SC, but also that of the resulting global change in transcriptional regulatory activity in the mutant strain, and these two contributions are difficult to distinguish. To avoid this issue, it is often preferred to use a wild-type strain, and induce a rapid SC variation by applying topoisomerase inhibiting antibiotics (8, 20). Commonly used drugs belong to the coumarin family (coumermycin, novobiocin), inhibiting the ATPase activity of gyrase (and topoIV), and the quinolone family (norfloxacin, ciprofloxacin, oxolinic acid) inhibiting the ligase activity of gyrase and topoIV (1, 3, 22). These drugs induce a sudden DNA relaxation in various species, whose effect on gene expression can then be measured. The main shortcoming is that they also trigger SC-independent stress-response pathways in the cell, which are then difficult to distinguish from SC-related transcriptional effects; this is especially true with quinolones, which induce double-strand breaks and a SOS response with pleiotropic effects. To minimise these issues, the drug concentration is usually kept as low as possible, and the expression is measured very quickly after the shock. Still, in order to characterise specifically the effect of SC on transcriptional regulation, it is thus highly desirable to increase the robustness of the analysis by comparing the expression patterns obtained with different methods (15). In this respect, a major limitation of existing studies is that, since gyrase is the primary target of all these drugs in Gram-negative bacteria, the transcriptomic response was analysed only in one direction, DNA relaxation, introducing a strong bias in the analysis of the SC-sensitivity of promoters.

The opposite variation could also be induced by applying quinolones on engineered strains harbouring mutant gyrase genes, where only the relaxing activity of topoIV is inhibited by the drug (2, 23). However, the steady-state SC level of these strains differs from that of wild-type ones, and the associated transcriptomic response was never monitored. In wild-type cells, topoI seemed a particularly suitable drug target (24), both in clinical research as it is the only enzyme of type IA topoisomerases family in many pathogenic species, but also as a way to study the effect of SC in transcriptional regulation, since this enzyme plays a direct role in the handling of torsional stress associated with transcription (25), while topoIV is predominantly involved in replication (26). Not only was it the first topoisomerase to be discovered (27), but the reduction or loss of its activity due to mutations in the *topA* gene in *E. coli* and *S. enterica* played an important role in the early discovery of the reciprocal interaction between transcription and SC (21, 28–30). In its catalytic cycle, topoI binds a stretch of single-stranded DNA, cleaves it and undergoes a conformational change to an open conformation, allowing the complementary DNA strand to pass the gate, followed by the religation of the DNA backbone with a gain of one linking number (31–34). In recent years, many compounds were shown to act as topoI inhibitors with unequal effectiveness as antimicrobial agents (24). In particular, one of them named seconeolitsine was shown to be effective against *Streptococcus pneumoniae* and *Mycobacterium tuberculosis* topoI, presumably by interacting with its nucleotide binding site, preventing the topoI conformational change and thus inhibiting DNA binding (34). When applied *in vivo* at low concentration, this drug induces a transient SC increase associated with a global change in the transcriptional landscape (35).

Here, we show this drug to be equally effective in Gram-negative bacteria, and we use it to report the first transcriptomic response to topoI inhibition and resulting SC increase in Gram-negative bacteria, using the phytopathogen *Dickeya dadantii* as a model. The latter contains the same set of topoisomerases as *E. coli* with a strong sequence homology, and generally, has a strong proximity to the enterobacterial models *E. coli* and *S. enterica*. But interestingly, SC was shown to be an important regulator of its key virulence genes (16, 36), and SC-affecting environmental signals are influential in its infection process, in particular osmolarity variations resulting in an increase of the cellular SC level (16, 36). Deciphering the mechanisms of SC-related transcriptional regulation in that species is thus important for our understanding of the mechanisms of virulence, as well as of transcriptional regulation as a general process.

In the following, we first demonstrate the inhibitory effect of seconeolitsine on *D. dadantii* (as well as *E. coli*) topoisomerase I, and its antibacterial action against that species. We then show that a seconeolitsine shock at lower concentration quickly increases the cellular SC level. We analyse the effect of this shock on the expression of the genome, and in particular, we illustrate the relationship between gene expression strength and spatial gene organisation and the response to topoI inhibition by seconeolitsine. By comparing this response with that of the gyrase inhibitor novobiocin and based on these contrasting observations, we propose a qualitative model explaining many notable features possibly involved in defining the supercoiling-sensitive property of promoters.

## Materials and Methods

### Seconeolitsine synthesis

Seconeolitsine was synthesised in 13% yield starting from boldine, following the protocol described in the original patent (37). In the first step, a reaction of demethylation was conducted in acidic conditions followed by a reaction with dibromomethane. The intermediate neolitsine was then reacted with chloroethyl chloroformate in dichloroethane followed by aromatization and ring opening achieved in refluxing methanol. The product was characterised by ESI-HRMS ([M+H]+: computed for C19H18NO4: 324.1230; found 324.1220) and its purity was validated by NMR (Supplementary Fig. S1).

### Protein expression and purification

*D. dadantii* 3937 *topA* gene was amplified and cloned into pQE80L plasmid using the TEDA method (38) to overproduce N-terminally 6xhis-tagged topoisomerase I. *E*.*coli* NM522 carrying the expression plasmid were grown at 37°C in LB medium until OD_600nm_ reached 0.6. Protein expression was then induced by adjusting the final concentration of the culture at 1 mM IPTG. After 2.5 h of induction, the cells were harvested by centrifugation, resuspended in a cold lysis buffer (20 mM NaH_2_PO_4_, 0.5 M NaCl, 20 mM imidazole, 2.5 mM TCEP (tris(2-carboxyethyl)phosphine), 1 mg/mL lysozyme, pH 7.4) (39) and disrupted through a French pressure cell press. After clarification of the obtained lysate by a 15 min centrifugation at 15000 rpm, the supernatants were mixed with Sigma HIS-Select Nickel Affinity Gel (at a ratio of 3:1) equilibrated in lysis buffer before being added into a polypropylene column (Qiagen). After extensive washing with a cold lysis buffer, the bound topoI was eluted with a cold elution buffer (20 mM, NaH_2_PO_4_, 0.5 M NaCl, 500 mM imidazole, 2.5 mM TCEP, pH 7.4) (39). Dialysis desalination was performed overnight with a first dialysis buffer (50 mM Tris-HCl, 100 mM NaCl, 0.1 mM EDTA, 1 mM DTT, pH 7.5) and 6 h with a storage buffer (50 mM Tris-HCl, 100 mM NaCl, 0.1 mM EDTA, 1 mM DTT, 50 % glycerol, 0.1 % Triton X-100, pH 7.5). The purity of topoI was assessed by SDS-PAGE and the concentration of the purified samples was measured with the Bradford protein assay (40). Comparisons with *E. coli* topoI were made with a commercial topoI (NEB).

### *In vitro* analysis of topoisomerase I inhibition by seconeolitsine

TopoI concentration required to relax 50% of pUC18 topoisomers was determined after 15 min of incubation at 37°C in rCutSmart Buffer (NEB). For the inhibition assays, TopoI was first preincubated with seconeolitsine and rCutSmart Buffer at 4°C for 10 min. This mix was then incubated with pUC18 at 37°C for 15 min. All reaction products were analysed by electrophoresis on 1.2 % agarose gel at 70 V for 3.5 h.

### Minimum inhibitory concentration (MIC) and survival rate in solid medium

LB Agar plates containing seconeolitsine dissolved in DMSO (50 mM stock solution) and IPTG (100 mM stock solution) were prepared to have seconeolitsine final concentrations between 0 and 750 µM and IPTG final concentrations of 0 or 0.1 mM. *D. dadantii* 3937, *E. coli* NM522 and *E. coli* NM522 carrying pQE80L::*topA* plasmids were grown at 30°C (*D. dadantii*) or 37°C (*E. coli*) until OD_600nm_ = 0.3. Cultures were then serial-diluted and placed on prepared plates. After 20 h of incubation at 30°C, colonies were counted. The survival rate was calculated as the ratio between the number of colonies observed on plates with or without seconeolitsine. The MIC was defined as the lowest seconeolitsine concentration without visible growth on the LB plates.

### Seconeolitsine inhibitory action in liquid cultures

*D. dadantii* 3937 were grown at 30°C in microplates containing Luria-Broth medium and increasing concentrations of seconeolitsine dissolved in DMSO (5 or 10 mM stock solution, keeping the final volume of DMSO below 4%). Optical densities were recorded every 5 min using an automatic microplate reader (Tecan Spark), and growth curves were fitted to a Gompertz equation to estimate growth rates and time lags (41).

### Bacterial cultures for seconeolitsine shock

*D. dadantii* 3937 were grown at 30°C in M63 supplemented with sucrose at 0.2% (wt/vol) until the exponential (OD_600nm_ = 0.2) or transition to stationary phase (OD_600nm_ = 1.1). Cells were then shocked with seconeolistine dissolved in DMSO at 50 µM during 5 min (RT-qPCR experiments) or 15 min (RT-qPCR and RNA-Seq experiments). An additional control was performed with pure DMSO for RT-qPCR experiments.

### Topoisomer separation in chloroquine-agarose gels

The topoisomer distribution was analysed as previously described (42). Reporter plasmids pUC18 were transformed into *D. dadantii* 3937. 15 minutes after the shock, plasmids were extracted with the Qiaprep Spin Miniprep kit and migrated on a 1% agarose gel containing 2.5 µg.ml^-1^ chloroquine at 2.5 V.cm^-1^ for 16 h. Under these conditions, more negatively supercoiled migrate faster in the gel. Chloroquine gels were subjected to densitometric analysis using Image Lab 6.0 software (Biorad). Distributions of topoisomers were normalised and quantified in each lane independently. IC50 was defined as the concentration that reduces topoI relaxing activity by 50%.

### RNA extraction

Total RNAs were extracted either with the frozen-phenol method (43) (RNA-Seq experiments) or with the Qiagen RNeasy Plus Mini Kit, including a bacterial lyse with a lysozyme solution at 1 mg.ml^-1^ and the optional DNase treatment (RT-qPCR experiments). The absence of genomic DNA contamination was further verified by PCR amplification with the Lucigen EconoTaq PLUS GREEN and *ryhB* primers (Tab. S1), following manufacturer’s instructions. When necessary, an additional DNase treatment was performed using the BioLabs DNase I to ensure RNA purity.

Extracted RNAs were quantified using a ND-1000 NanoDrop spectrophotometer. RNA quality was checked by agarose gel electrophoresis.

### Quantitative real time PCR

1 µg of total RNAs were reverse transcribed using the Thermo Scientific RevertAid First Strand cDNA Synthesis Kit. Reaction mixes were incubated at 25°C for 5 min, 42°C for 60 min and 70°C for 5 min.

The quantitative PCR was carried out using the Thermo Scientific Maxima SYBR Green/ROX qPCR Master Mix with the LC480 Lightcycler from Roche and the primers listed in Tab. S1. The following thermal cycling reactions were executed: (i) an initial denaturation step at 95°C for 10 min, (ii) 45 amplification cycles at 95°C for 15 s, 58°C for 30 s and 72°C for 40 s. The housekeeping gene *rpoA* was used as a normalizer for the gene expression ratios. The uniqueness of the amplification product is verified with the melting curve.

### RNA Sequencing

All samples were collected in two biological replicates (8 samples in total). Steps of ribosomal RNA depletion, cDNA library preparation and high-throughput sequencing were carried by the MGX Montpellier GenomiX platform, using the Illumina TruSeq stranded mRNA sample preparation kit and HiSeq2500 sequencing providing 50-nt single-end reads. The sequenced reads were deposited in ArrayExpress under accession number E-MTAB-10134. They were mapped on the reference genome of *D. dadantii* 3937 (NCBI NC_014500.1) with Bowtie2 and counted with htseq-count. Gene differential expression analysis was performed with DESeq2 with a threshold of 0.05 on the adjusted p-value.

### Statistics and data analysis

All statistical analyses and graphs were made with a homemade Python code. Error bars are 95% confidence intervals. Proportions of activated genes among differentially expressed genes were compared with χ^2^-tests. Stars indicate the level of significance based on the p-value (***, *P <* 0.001; **, 0.001<*P*<0.01; *, 0.01<*P*<0.05). The orientation of a gene is defined relative to the orientation of its neighbours (either convergent, divergent or tandem). Functional enrichment was analysed using the Gene Ontology classification (44). Only functions corresponding to at least four *D. dadantii* genes were considered. Chromosomal domains were previously defined in (16).

## Results

### Seconeolitsine inhibits *D. dadantii* topoisomerase I *in vitro*

The comparison of *topA* sequences from enterobacteria *D. dadantii* and *E. coli* with those of *M. tuberculosis* and *S. pneumoniae* showed that the topoI residues bound by seconeolitsine were mostly conserved in the former (Supplementary Fig. S2), suggesting that the inhibitory activity of the drug might be also effective in enterobacteria. To test this hypothesis, we synthesised seconeolitsine, following the protocol described in the original patent (37), and the purity of the product was validated by NMR (Supplementary Fig. S1). The inhibitory activity of econeolitsine against *D. dadantii* topoI was evaluated by adding increasing concentrations of the drug to a solution of purified enzymes (see Materials and Methods), resulting in a progressive reduction of their relaxing activity with a IC50 of 4.6 μM (Fig. 1), similar to that observed with *M. tuberculosis* topoI (45). We also observed an inhibitory effect on purified topoI from *E. coli*, with a IC50 of 6.7 µM (Supplementary Fig. S3). Together, these observations also suggest that seconeolitsine might be effective against topoI from a broader variety of bacterial species. Note that topoI from *D. dadantii* had a stronger apparent relaxing activity than that of *E. coli* (Supplementary Fig. S3) and was therefore added in lower concentration in the assay, which might contribute to the quantitative difference in IC50 between these two species, but this question requires further investigation.

**Fig. 1:**
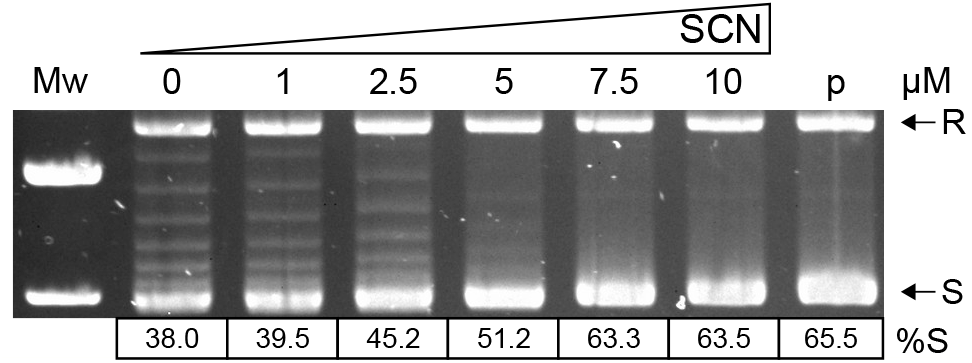
Inhibition of topoI *in vitro* relaxing activity by seconeolitsine. The specified amount of seconeolitsine was pre-incubated for 10 min with 100 ng topoisomerase I, then 0.5 µg of supercoiled pUC18 plasmids (p) was added and incubated for 15 min. %S is the percentage of topoisomers in the highly negatively supercoiled form (S). R indicates the relaxed form.

### High concentrations of seconeolitsine impede *D. dadantii* growth

We then investigated the antibacterial activity of the drug, by analysing its effect on *D. dadantii* growth. In solid medium, we observed a progressive reduction in bacterial growth, with a minimal inhibitory concentration (MIC) of around 500 μM (Fig. 2A). In liquid cultures in microplates (Fig. 2B), we observed that the drug increasingly impedes growth, with a lag time proportional to the applied dose in the 100-300 μM concentration range. Altogether, the antibacterial effect of the drug occurs at much higher concentrations in *D. dadantii* than *M. tuberculosis* (MIC of 500 μM versus 16 μM). Since the *in vitro* IC50 values are comparable for the topoI enzymes from the two species, this strong difference presumably arises from cellular properties (in particular the membrane structures), resulting in a different bioavailability of the drug molecules in the cells. As a comparison, *E. coli* cells were inhibited by lower concentrations of seconeolitsine than *D. dadantii*, with a MIC of around 250 μM (Supplementary Fig. S4), whereas the growth of the Gram-positive bacterium *Bacillus subtilis* is impeded already at concentrations around 20 μM, comparable to those of *S. pneumoniae* (Supplementary Fig. S5).

**Fig. 2:**
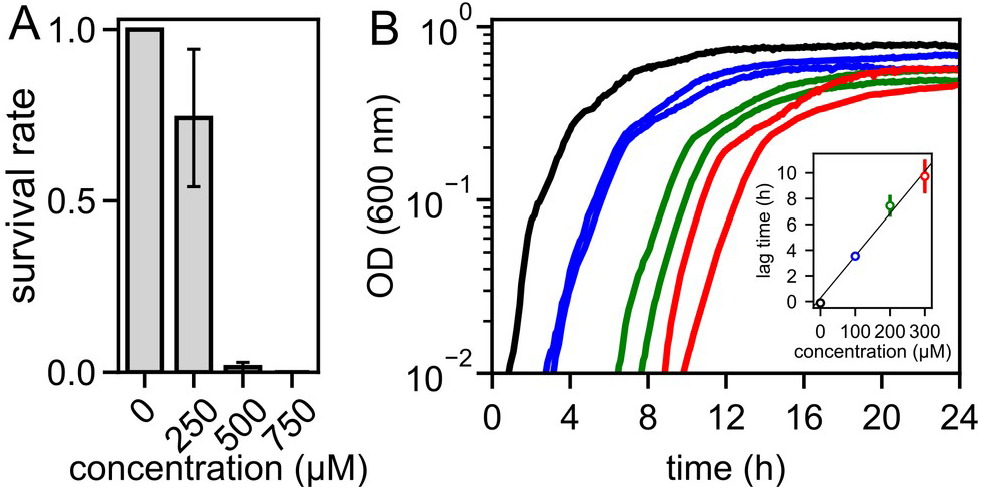
Antibiotic effect of seconeolitsine on *D. dadantii* 3937. (A) Survival rate in the presence of increasing amounts of seconeolitsine in solid medium. Each bar indicates the proportion of growing colonies with the specified amount of seconeolitsine in comparison to plates without seconeolitsine, with a 95% statistical confidence interval (see Materials and Methods). (B) Growth curves in the presence of increasing amounts of seconeolitsine in liquid medium. The linear increase of the lag time with drug concentration, obtained from a quantitative analysis of growth curves (see Materials and Methods) is shown in the inset.

In order to confirm that *D. dadantii* topoI is indeed targeted by seconeolitsine *in vivo*, we analysed the effect of overexpressing the enzyme on cell survival, in a medium containing the drug at a partially inhibitory concentration (Supplementary Fig. S6). While the survival rate is around 30% in absence of the inducer, it is significantly higher (73%, *P*=0.0015) when topoI is overexpressed. This observation suggests that at least a significant fraction of the seconeolitsine molecules are indeed targeted to topoI, although other cellular effects cannot be excluded.

### Seconeolitsine shock increases DNA superhelicity in *D. dadantii* cells

Based on the previous observations and in line with previous studies (35), we anticipated that a seconeolitsine shock at a sublethal concentration might transiently inhibit the activity of topoI in *D. dadantii* cells and thus induce a rapid SC increase. Indeed, a concentration of 50 μM induced a significant shift in the distribution of topoisomers of the pUC18 plasmid extracted 15 min after the shock (Fig. 3A, this time delay was previously chosen to monitor the impact of novobiocin in *D. dadantii*). This concentration was used in all further experiments, because at the same time, it was sufficiently weak to avoid any observable effect on the growth of exponentially growing cells (Supplementary Fig. S7), thus minimising general physiological side-effects of the shock versus the direct regulatory effect of DNA SC that we wish to investigate.

**Fig. 3:**
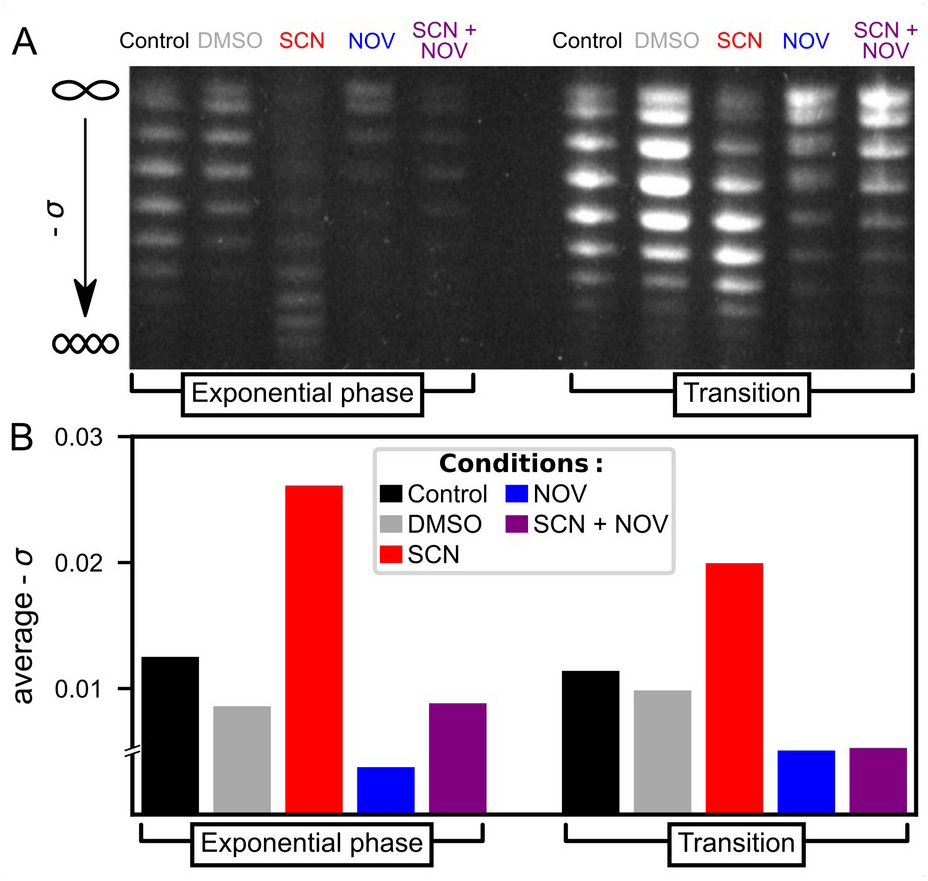
A seconeolitsine (SCN) shock at 50 μM concentration induces an increase of SC in cellular DNA after 15 min. Conversely, a novobiocin (NOV) shock at 100 µg.ml^-1^ induces DNA relaxation. (A) Agarose-chloroquine gels of pUC18 plasmids isolated from *D. dadantii* 3937 cells. At the employed concentration of chloroquine, the downward migration increases with SC level, and the SC increase induced by seconeolitsine can be fully resolved. (B) Average negative SC level computed from the quantification of the topoisomer distribution. Note that the most relaxed fraction of the topoisomer distribution in presence of novobiocin was not fully resolved, preventing an exact estimation of the relaxation magnitude in these samples, hence the discontinuity indicated in the y-axis.

The distribution of topoisomers is entirely resolved in the untreated and seconeolitsine-treated samples, and thus allows an unambiguous quantification of the observed profiles. In the treated cells, the average negative SC level is increased by Δσ=-0.014 in exponential phase and Δσ=-0.009 at the transition to stationary phase (quantified topoisomer distributions are available in Supplementary Fig. S8). The weaker effect observed at the latter stage was expected since both gyrase and topoI are more active in the exponential phase (8, 10, 46). We checked that this increase is absent when only DMSO is applied. In both phases, the sharp increase in SC induced by seconeolitsine is in clear opposition to the relaxed levels measured after novobiocin treatment (36), as we expected based on the opposite activity of topoI vs gyrase. Accordingly, in a control sample where both drugs are added simultaneously (rightwards lanes), the plasmids reach an intermediate superhelical level.

Since there are no previous studies of topoI inhibition in *D. dadantii*, these data cannot be directly compared to previously published data; however, the shift in topoisomer distributions observed after seconeolitsine treatment is qualitatively similar to that observed after an osmotic shock (36), which is also known to increase the negative superhelical level in *E. coli, S. typhimurium* and several other species (8). An additional experiment shows a similar effect in *E. coli* cells, albeit with a stronger effect of seconeolitsine at this concentration of 50 μM (Supplementary Fig. S9).

### Transcriptional response of selected promoters

We expected the global increase in negative SC level to affect the expression of many genes of the *D. dadantii* chromosome, and therefore analysed the transcriptional effect of the seconeolitsine shock using RNA-Seq, with a qRT-PCR validation of selected genes. We first illustrate the kinetics of the transcriptional response of four genes strongly responsive to seconeolitisne: the *dps* gene encoding the NAP Dps, which is possibly the most abundant DNA-binding protein in stationary phase (47) and condenses the chromosome under conditions of resource scarcity or stress; the *desA* gene involved in efflux systems; a gene of unknown function (accession number *Dda3937_02096*); and *feoA* involved in iron transport. In the exponential phase (Fig. 4A), these genes react very quickly (5 min) and in opposite manners. The response measured by RNA-Seq after 15 min (B) was entirely consistent with that measured by qRT-PCR (A); in the latter, we confirmed that DMSO (used as solvent) triggers no detectable transcriptional response (thin lines), indicating that seconeolitsine is indeed the active molecule. Similar effects were observed at the transition to stationary phase (Fig. 4 C and D). The functions of these strongly responsive genes suggest that they are part of a mechanism of drug-response by the bacteria, which is not surprising given the antibacterial nature of the drug. On the other hand, SC modulates the expression of many genes in a global but usually milder manner (48), as can be observed in Fig. 5 with genes expected to respond specifically to SC variations.

**Fig. 4:**
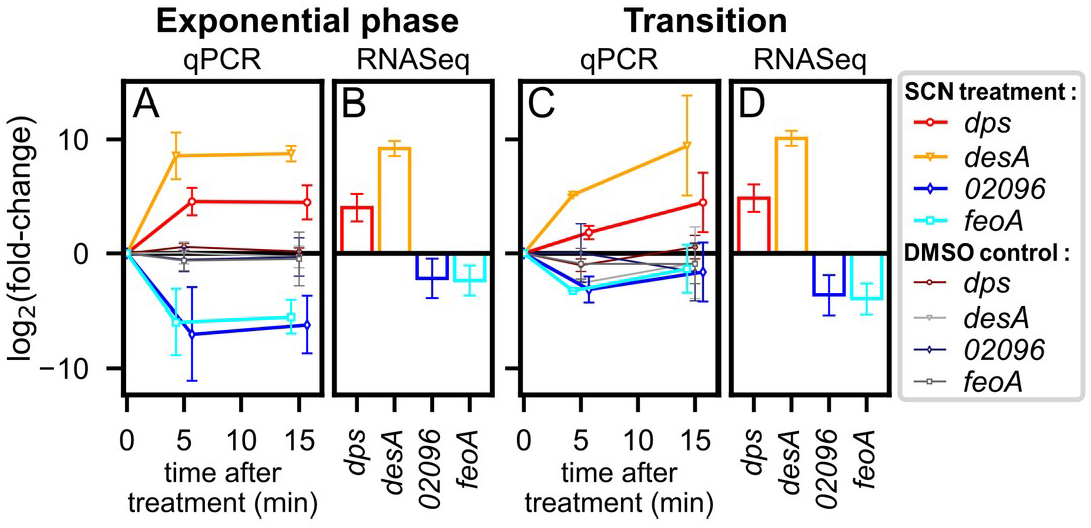
Kinetics of promoter activation (*dps, desA*) or repression (*Dda3937_02096, feoA*) by seconeolitsine (SCN) shock. Gene expression levels were measured in exponential phase (A and B) and at the transition to stationary phase (in C and D), either by qRT-PCR (5 and 15 min post-shock, coloured markers and thick lines in A and C) or by RNA-Seq (after 15 min incubation with seconeolitsine, B and D). Control datapoints obtained after incubation with the same volume of pure DMSO solvent are indicated as thin lines, and exhibit no detectable effect. All error bars shown indicate 95% confidence intervals, obtained with 3 biological replicates (qRT-PCR) or from RNA-Seq analysis.

**Fig. 5:**
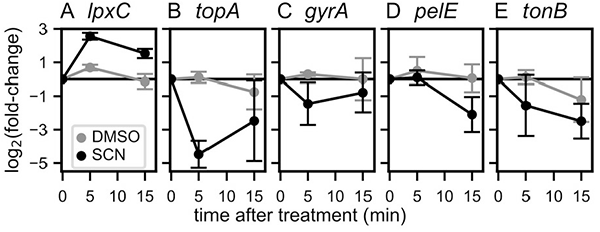
Effect of seconeolitsine shock on various genes expected to respond to variations of SC: (A) *lpxC*; (B) *topA*; (C) *gyrA*; (D) *pelE;* (E) *tonB*. The expression was measured by qPCR 5 and 15 min after the shock in the exponential phase (black dots). Control datapoints treated with the DMSO solvent are shown (grey dots).

The *lpxC* gene illustrates some difficulties encountered when analysing SC-controlled regulation. That gene was previously identified as particularly stable in the presence of various changes of environmental conditions, and because of this apparent lack of regulation, was considered as a suitable internal normalizer for qRT-PCR experiments with *D. dadantii* (49). However, *lpxC* was later found to be sensitive to DNA relaxation by novobiocin (16, 50). Similarly here, its expression is increased by seconeolitsine treatment in both growth phases, as observed in both qRT-PCR experiments (Fig. 5A) and RNA-Seq data (Supplementary Tab. S2), the former being either calibrated by concentration gradient or using *rpoA* as internal normalizer. This example shows that SC may affect the expression of a large class of promoters, possibly even those lacking a direct dependence on transcription factors since it modulates the direct interaction of RNAP with promoter DNA (8). The expression of *rpoA*, on the other hand, appeared stable in the investigated conditions and it was used as a normalizer for all qRT-PCR data presented.

We then investigated the response of topoisomerase genes. The *topA* gene was found to be repressed by the shock (Fig. 5B, not significant in the less sensitive RNA-Seq data), in agreement with observations in *S. pneumoniae* (35). The authors of the latter study analysed the kinetics of SC homeostasis, with a strong initial SC increase followed by a partial relaxation. However, it must be noted that this repression of the *topA* gene in both species contradicts the behaviour expected for a simple homeostasis mechanism, which would lead to an activation, comparable to the homeostatic activation of the *gyrA/B* genes by gyrase inhibition (4). In a different study, conversely, the *topA* promoter was activated by an increase in negative SC induced by oxolinic acid in mutant *E. coli* cells (51), suggesting that its response is possibly more versatile and condition-dependent than that of *gyrA/B* genes. Since the basal SC level is more relaxed in the employed *E. coli* strain (gyrase mutant), and the magnitude of SC variation is weaker, a possible explanation is that the very high negative SC level reached after seconeolitsine treatment might exceed the dynamic range of the homeostatic response of the *topA* promoter.

Among other topoisomerases, we observed a slight repression of *gyrA* expression (Fig. 5C, not significant in the RNA-Seq data) as well as a possible activation of gyrase inhibitors (the *Dda3937_01484* gene, associated to this function by sequence homology, was found significantly activated in the RNA-Seq data, but not confirmed by qRT-PCR). No effect on topoIV genes (*parC/E*) was detected. Altogether, the regulatory mechanisms of SC homeostasis in response to topoI inhibition remain to be fully characterised, and might thus involve a rapid reduction of gyrase activity in addition to changes in topoI expression.

We looked at the *pelE* gene, which encodes a major virulence factor of *D. dadantii* responsible for plant cell wall degrading activity, and is strongly repressed by novobiocin (36). *pelE* was repressed by seconeolitsine in exponential phase (Fig. 5D), and not significantly affected at the transition where topoI activity is weaker. The fact that this gene is repressed by both novobiocin (relaxation) and seconeolitsine (SC increase) suggests that the expression is optimal at the natural SC level, consistent with the tight regulation of this level in the cell (21).

Finally, we investigated the *tonB* gene, involved in iron siderophores and vitamin B_12_ transport at the cell membrane. This gene was previously found to be repressed by an increase of SC induced by anaerobiosis in both *E. coli* and *S. enterica*, and this repression was relieved by a novobiocin treatment restoring a SC level close to the physiological one (52). Similarly here, we observed a strong repression of *tonB* expression by seconeolitsine in both qRT-PCR experiments (Fig. 5E) and RNA-Seq data (Supplementary Tab. S3) in the exponential phase, giving further support to the repressive effect of strongly negative SC levels on that promoter.

### Global transcriptional effect of seconeolitsine shock

A comparison in the response of several genes confirms that qRT-PCR and RNA-Seq results are well correlated (Supplementary Fig. S10), leading us to analyse the transcriptomic results at the global scale. Such an analysis is particularly useful for SC-related regulation, which affects many genes in a quantitatively mild manner, because it can unravel global regulatory patterns undetectable at the level of individual promoters (16, 35). While most previous studies of gyrase inhibition in Gram-negative bacteria were carried in the exponential phase only (14–16, 19), we have measured the response to seconeolitsine treatment in the two stages of growth, in each case 15 min after the shock. The lists of differentially expressed genes in either growth phase are given in Supplementary Tables S2 and S3).

The distributions of affected genes are provided in Fig. 6. The shock has a significant impact on around 13% of the genome in the exponential phase, and 7% at the transition to stationary phase. Remarkably, only a small minority of the genes respond significantly in both phases, in which case the response goes in the same direction, whereas many genes respond significantly only in one phase (e.g., *tonB*). This behaviour was not unexpected, since the chromosome conformation (including SC level) and topoisomerase activities are quite different in exponential phase vs transition to stationary phase (10, 46). Among differentially expressed genes, the proportion of activated vs repressed ones is considerably higher at the transition than in the exponential phase (Fig. 6C).

**Fig. 6:**
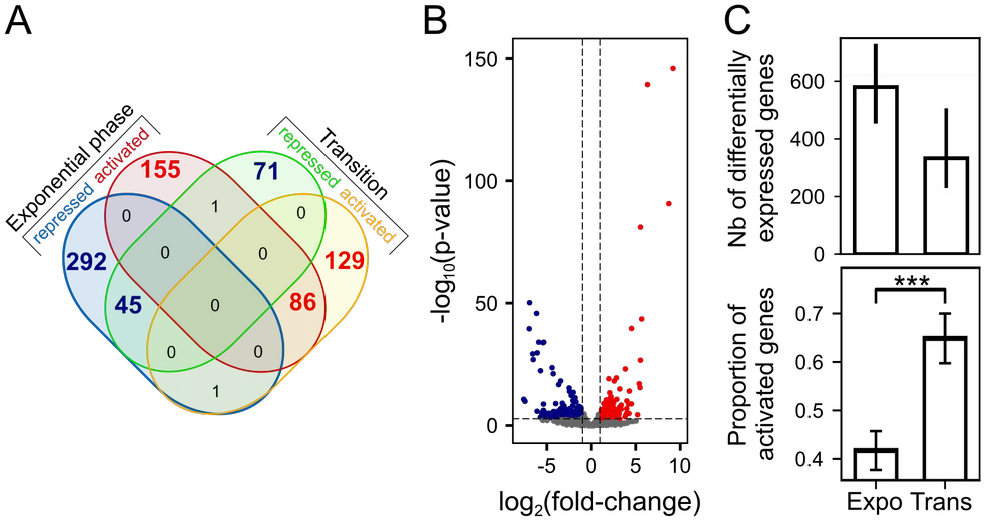
Global response of the *D. dadantii* genomic expression to a seconeolitsine shock. (A) Venn diagram of significantly activated or repressed genes in the two growth phases, with a threshold of 0.05 on the adjusted p-value. The red numbers refer to genes activated by seconeolitsine (in either phase), and the blue numbers to repressed genes. These numbers vary by around 30% when the threshold is changed by a factor 2. (B) Volcano-plot showing the genomic response to a seconeolitsine shock in exponential growth. Red dots and blue dots correspond to activated and repressed genes, respectively (p-value threshold of 0.05 and |log2(fold-change)| threshold of 1 are indicated as dashed lines). Unaffected genes are shown in grey. (C) Top: Total number of differentially expressed genes (among 4260 genes in total) with an adjusted p-value threshold of 0.05 (bars indicate variations of this number with thresholds of 0.025 and 0.1). Bottom: proportion of activated genes among them (with 95% statistical confidence intervals).

In previous studies, DNA relaxation was shown to regulate the expression of the genome in a functionally scattered way, with limited enrichment in specific regulatory pathways (48). We therefore analysed if the same is true of the seconeolitsine shock (Fig. 7). Indeed, relatively few Gene Ontology (GO) categories exhibit a strong systematic response, and they belong to very diverse functional groups. Expectedly, the most present pathways are related to (i) metabolism and biosynthesis, as already observed during DNA relaxation (14), which are affected differently in the two phases (see grey, blue and red groups in Fig. 7); and (ii) transport and efflux systems, which may, in part, participate in the cellular response to the drug, and are mostly affected similarly in the two phases (green group in Fig. 7). We also noted a strong activation of the iron metabolism pathway. But importantly, these enriched functions comprise less than 40% of the total number of differentially expressed genes, showing that most genes are regulated separately rather than within their entire functional category. This feature is characteristic of the “analogue” regulation mode by global changes of the chromosome conformation (48), which affects genes according to their spatial localisation and organisation, in contrast to the classical “digital” regulation by transcriptional regulatory networks, where each regulator often binds many promoters within a functional group. We now look in more detail at such spatial organisational features of the global pattern of expression.

**Fig. 7:**
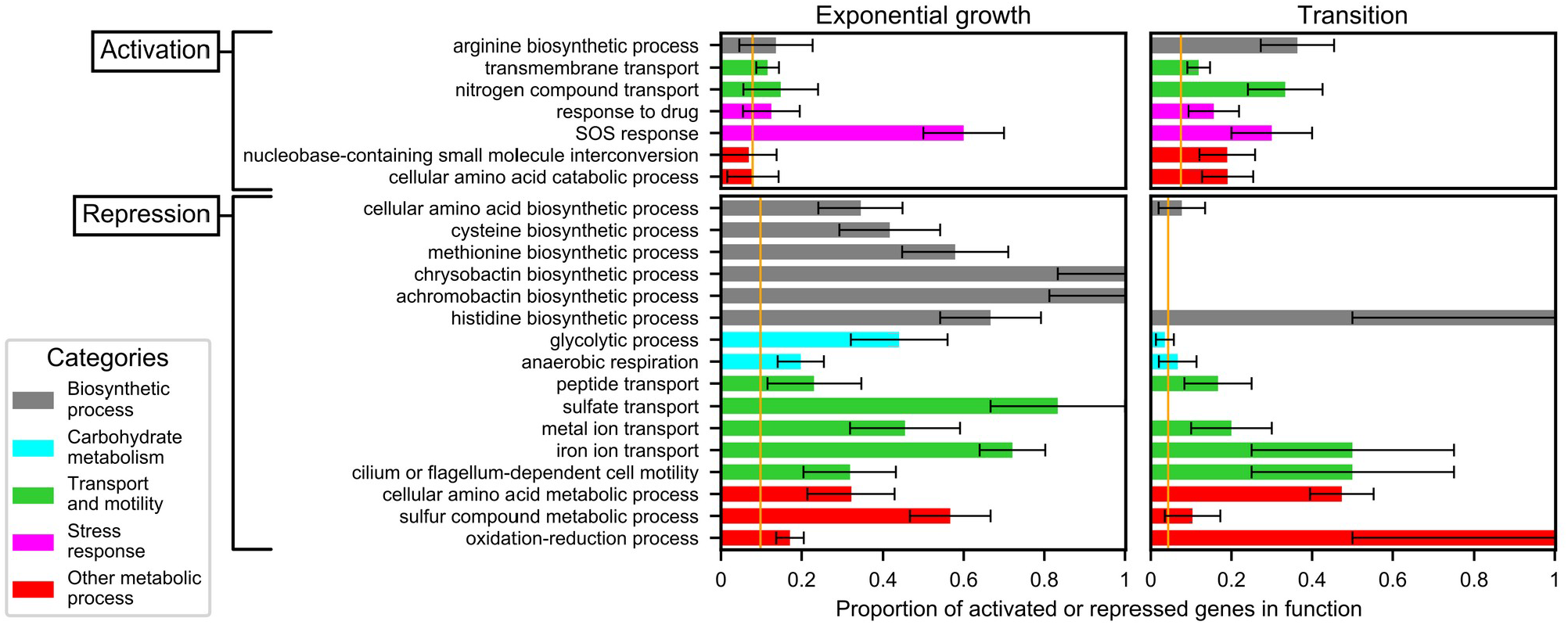
Functional enrichment analysis of activated (top) or repressed (bottom) genes, during a shock in exponential (left) or transition to stationary phase (right). Each bar indicates the proportion of differentially expressed genes in the considered function (with a 95% statistical confidence interval), which can be compared to the genomic average (orange vertical lines): the considered function is enriched if the confidence interval does not cross the orange line. Colours indicate the repartition in broad functional groups.

### Spatial organisation of promoters sensitive to seconeolitsine shock

We started by representing the large-scale distribution of regions enriched in activated or repressed genes along the chromosome (Fig. 8). Strikingly, whereas these regions are almost identical in the two investigated growth phases during a novobiocin shock (Fig. 8B), they are essentially different during a seconeolitsine shock (Fig. 8A), suggesting that, while the large-scale distribution of gyrase activity is similar in the two growth phases, that of topoI is growth phase-dependent.

**Fig. 8:**
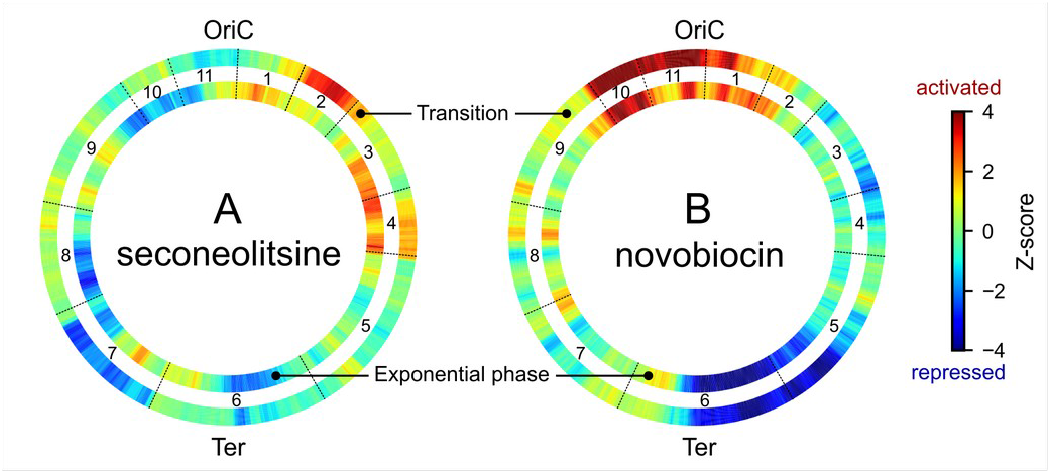
Distribution of genomic regions enriched in activated (red) or repressed (blue) genes, in exponential phase (internal wheels) or transition to stationary phase (external wheels), during topoI inhibition by seconeolitsine (A) or gyrase inhibition by novobiocin (B). The colours represent the statistical significance of the proportion of activated over repressed genes in sliding 500-kb windows (Z-score > 2 or < -2, respectively); if the number of differentially expressed genes in the window is low, the Z-score remains close to 0 and appears in green. 11 domains of coherent expression (CODOs) previously identified (16) are indicated.

Previous analyses of *D. dadantii* transcriptomes led to the definition of eleven domains of coherent stress-response, termed CODOs (16, 52), which harbour distinct DNA physical properties, are differentially regulated by NAPs and novobiocin, and respond coherently to various stress signals encountered during plant infection. These domains are indicated in Fig. 8 (black boundaries between the wheels), and in many cases, coincide with patterns of topoI activation/repression. As an example, domain 7 (bottom left) harbouring several virulence genes (type VI secretion systems, flagella and chemotaxis operons) is repressed by topoI inhibition at the transition to stationary phase. Interestingly, the same effect is observed when the bacteria are subjected to an osmotic shock at this stage of growth (16, 52), which also triggers an increase in negative SC (16, 36), and mimics the physiological conditions encountered at the beginning of the maceration phase of plant infection (53). Other domains are repressed in exponential phase (domain 10), or activated either in exponential phase (domain 4) or at the transition (domain 2). Again, this latter observation is consistent with the effect of an osmotic shock, which down-regulates catabolic activity in general and specific stress-responsive genes in domain 2 in particular (16). In summary, although the physical nature and the mechanisms underlying the emergence of these domains remain to be clarified, the transcriptional effect of seconeolitsine gives further support to the notion that they reflect an architectural ordering of the chromosome involving SC and affecting its expression, in line with comparable observations in *S. pneumoniae* (54).

### Topoisomerase I inhibition hinders the expression of strong promoters

A notable feature of the large-scale expression pattern (Fig. 8) is that, while the gyrase inhibition pattern is characterised by a clear ori/ter vertical asymmetry (B), the topoI inhibition pattern rather displays an approximate left/right replichore asymmetry (A). However, a statistical comparison of the proportions of activated genes did not exhibit any global difference between the left and right replichores, suggesting that this difference is rather localised in specific regions. Rather, we did find a higher proportion of activated genes on the lagging vs leading strand on transition to stationary phase (Supplementary Fig. S11), suggesting that topoI is more important to dissipate negative supercoils on the leading strand (considering both replichores, i.e., with RNAP and DNA polymerase translocating either in the same or in opposite directions). A putative explanation is that the topological constraints might be weaker on the lagging strand where replication does not proceed continuously, but since this difference is not observed in exponential phase where replication is more active, it is likely that the topological constraints induced by the latter are then efficiently handled by topoIV.

Since the leading strand is known to be enriched in highly expressed genes, we looked for a relationship between expression strength and response to seconeolitsine treatment. TopoI is known to colocalise with RNAP and possibly release negative supercoils in its wake at strongly expressed promoters (25), and we therefore expected the latter to be particularly hampered by topoI inhibition. Indeed, we found a very strong and progressive increase in the proportion of inhibited genes depending on their expression level in the exponential phase (Fig. 9A), from 23% (of all differentially expressed genes) in the lowest quartile, up to 65% in the highest quartile. In line with previous results (25), this observation suggests that topoI participates in active transcription, not only at a few highly expressed operons, but as a global mechanism. In contrast, at the transition to stationary phase, the proportion of inhibited vs activated promoters was almost independent of promoter strength (Fig. 9B; note that strongly expressed promoters were more responsive, but in similar proportions in both directions, Supplementary Fig. S12). Possible reasons include a globally weaker transcription level at the transition, or a weaker inhibitory effect of the transcription-induced negative supercoils in the latter phase where the global SC level is more relaxed (see Discussion).

**Fig. 9:**
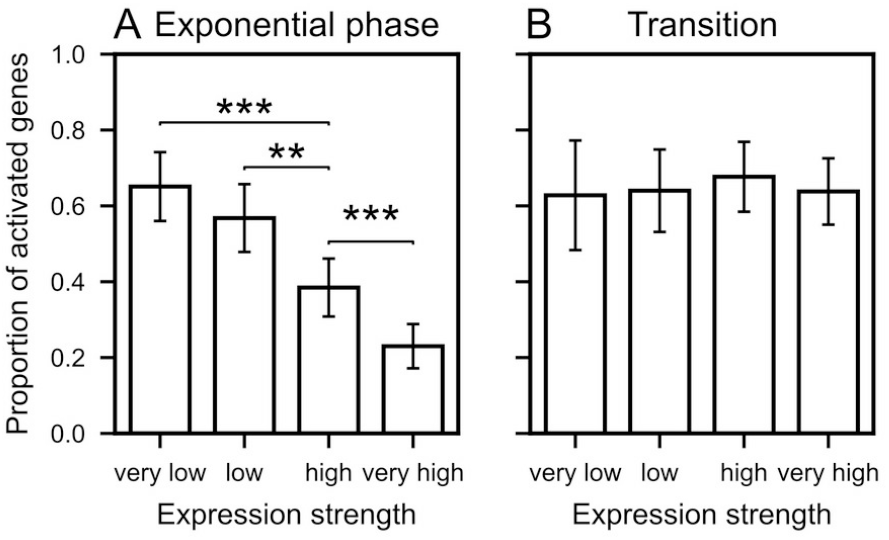
Proportion of activated genes among differentially expressed genes depending on expression strength, in the exponential phase (A) or at the transition to stationary phase (B). Error bars indicate 95% confidence intervals, and significant differences are indicated (no effect at the transition). Genes were separated into quartiles based on their average number of reads in all samples. The proportions of differentially expressed genes are shown in Supplementary Fig. S12.

### Role of neighbouring gene orientation

We then investigated a possible relation between neighbouring gene orientations and the response to seconeolitsine. Such a relationship was expected for the same reason as the previous observation, since RNAP-generated supercoils accumulate not only behind actively transcribed genes (25), but more specifically between divergent operons (55), as occurs at the *leu* promoter of *S. enterica* where this mechanism was discovered (30, 56). The orientation of a gene is here defined by the coding DNA strands of its two neighbours relative to it (in the case of tandem genes, the two neighbours belong to the same strand, which can either be the same as the considered gene or the opposite one). Fig. 10 shows that the expected dependence is indeed observed in both growth phases, with genes located between divergent neighbours being significantly more repressed by topoI inhibition compared to convergent ones. This observation, made at the scale of the entire genome, shows that the regulatory mechanism uncovered at the *leu* locus (57) is widespread in the *D. dadantii* genome (and likely in other Gram-negative bacteria). A similar effect of gene orientation had been also observed following novobiocin treatment (Supplementary Fig. S13), highlighting the tight relationship between topoisomerase activity and the genomic organisation due to transcription-induced supercoils (50).

**Fig. 10:**
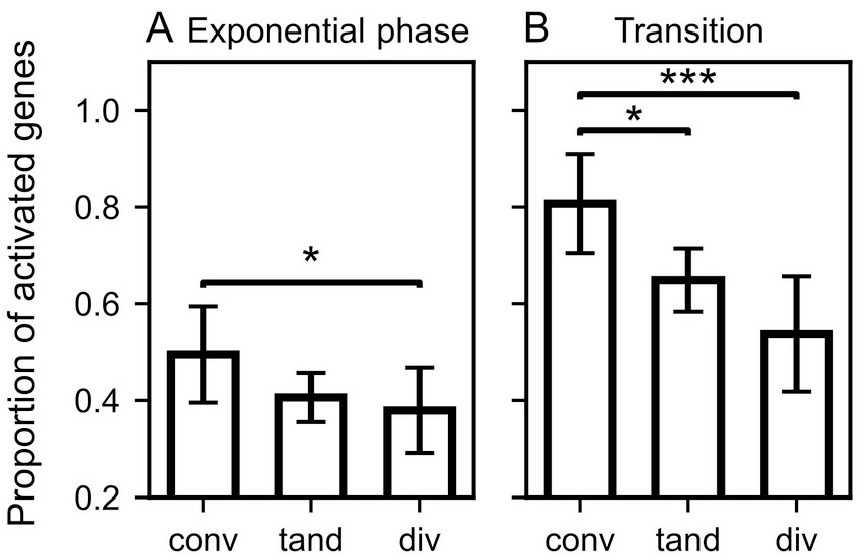
Gene orientation-dependent transcriptional response to seconeolitsine. The proportion of activated genes (among differentially expressed ones) is significantly higher among those located between convergent (conv) than divergent (div) neighbours, both in exponential phase (A) and at the transition to stationary phase (B). Tandem (tand) genes exhibit intermediate values. Number of differentially expressed genes (conv, tand, div): 57, 205 and 67 in exponential phase, and 97, 362 and 116 at the transition, respectively.

## Discussion

### The supercoiling-sensitivity of promoters is condition-dependent

We have collected the first transcriptomic response to an increase of SC due to topoI inhibition in a Gram-negative bacterium. Since all previous analyses in these species involved the opposite variation, DNA relaxation induced by gyrase inhibition, we wished to compare these complementary responses, in order to refine our understanding of the notion often referred to as the “supercoiling-sensitivity” of promoters.

Fig. 11 shows that, among genes responding to one of the drugs, the large majority does not respond to the other: genes appearing as sensitive to DNA relaxation are therefore essentially different from those sensitive to an increase of SC. This observation is possibly affected by the limited sensitivity of the RNA-Seq experiment, where some genes confirmed by qRT-PCR (*pelE, gyrA*) fell below the threshold of statistical significance. Among the genes responding to both drugs, most of them do in the same direction, including some belonging to stress-response functions of the cells possibly via SC-independent regulatory pathways (such as *dps* or *desA*) but also some likely directly regulated by SC (such as *pelE, topA* or *tonB*). Finally, a remarkably low number of genes respond in opposite directions to the two drugs, as would yet be naively expected from promoters exhibiting an intrinsic and general property of supercoiling-sensitivity. Note that the latter proportions of similar vs opposed responses to the two drugs were quite comparable in *S. pneumoniae* cells in exponential phase (Supplementary Fig. S14).

**Fig. 11:**
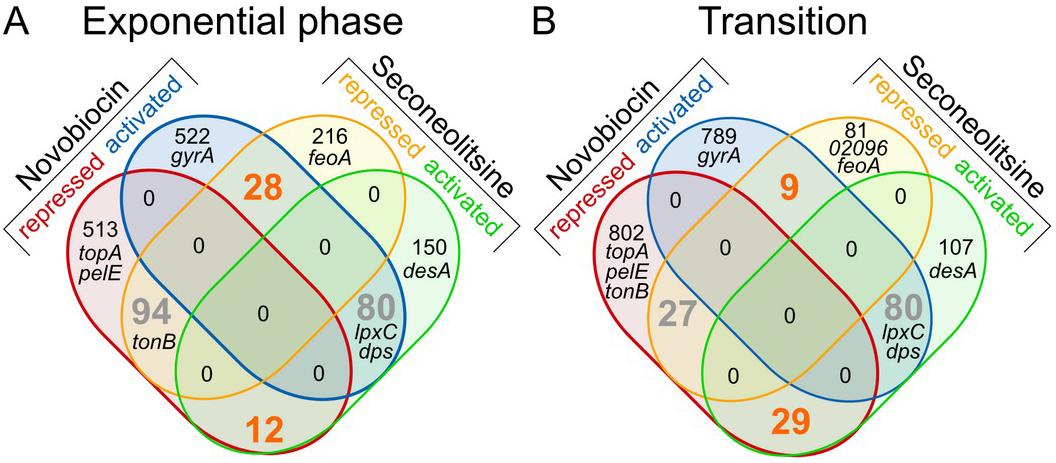
Venn diagrams of genomic response to novobiocin and seconeolitsine, in either exponential phase (A) or transition to stationary phase (B). The number of genes responding in opposite directions to the two drugs are indicated in orange, and those in the same direction in grey. Selected genes are indicated in their respective categories. 02096 is the gene with accession number *Dda3937_02096*.

These observations, together with others made in this study, highlight the complexity of the SC-related regulation of transcription. The response of a given promoter depends on the action of global parameters related to the physiology of the cell (growth phase, metabolic state, …) but also to more localised and dynamic factors that are difficult to decipher (local activity of topoisomerase enzymes, mechanical effects of local transcription and replication, binding of nucleoid-associated proteins, …), explaining the current lack of predictive and quantitative models of this form of regulation, as opposed to those involving transcriptional factors (58).

### A qualitative model for the response of bacterial promoters to global variations of DNA supercoiling

In spite of these difficulties, is it possible to propose at least a qualitative model of transcriptional regulation, depending only on the average SC level measured with agarose-chloroquine gels as above, and recapitulating the most notable features observed in transcriptomes obtained with both gyrase and topoI inhibitors?

In order to focus on the most generic features of SC-dependent transcriptional regulation, it is useful to analyse *in vitro* transcription data, where genes are expressed on plasmids with minimal influence of genomic context or regulatory proteins, and where SC is modified in a controlled way. Fig. 12A recapitulates several available datasets of this kind (7, 11, 59) obtained with a broad sampling of SC levels comprising typical physiological levels, either in standard conditions (from – 0.04 to -0.06), upon gyrase inhibition (lower negative SC levels) and upon topoI inhibition (higher negative SC levels). The employed promoters belong to different promoter families, either from stable RNAs (*tyrT*) and mutant promoters derived thereof (*tyrTd*), or promoters of protein-encoding genes (*galP*) or derivatives of *lacP* (*lacPs, lacPsd*). And yet apart from conspicuous differences between these curves, a similar pattern is clearly and repeatedly observable: the expression is very low on an entirely relaxed DNA template, then increases drastically and monotonously until reaching maximal expression at a (promoter-dependent) optimal SC level close to the physiological level in exponential phase (≈ -0.06), then decreases at higher SC levels. This behaviour is schematised in Fig. 12B, where the horizontal axis is voluntarily left without quantitative values. The two background colours highlight the two regulation regimes with putative associated mechanisms: the initial activation curve is likely due to the SC-induced reduction of DNA opening free energy during open-complex formation, which occurs preferentially at the highly AT-rich region starting at the -10 promoter element where the transcription bubble is formed (17); the decrease is more complex and either due to the opening of secondary sites competing with the -10 element (60), or to a reduction in processive initiation due to an excessive stability of the open-complex resulting in more abortive transcripts (6). Accordingly, while a modification of the AT-richness of the promoter sequence downstream of the -10 element (in *lacPsd* and *tyrTd* compared to *lacPs* and *tyrT*, respectively) clearly shifts the activation curve horizontally in a predictable manner (17), the second part of the curve is more variable, and the position of the maximum differs significantly from one promoter to the other (7). Based on this empirical model, the regulatory effect of a SC variation (e.g., due to a topoisomerase inhibitor) is then expected to depend both on the initial global SC level in the cell, and on the direction and magnitude of the SC change. Approximate values of the average SC level in exponential or transition to stationary phase are indicated in blue, with exact values varying between species (61).

**Fig. 12:**
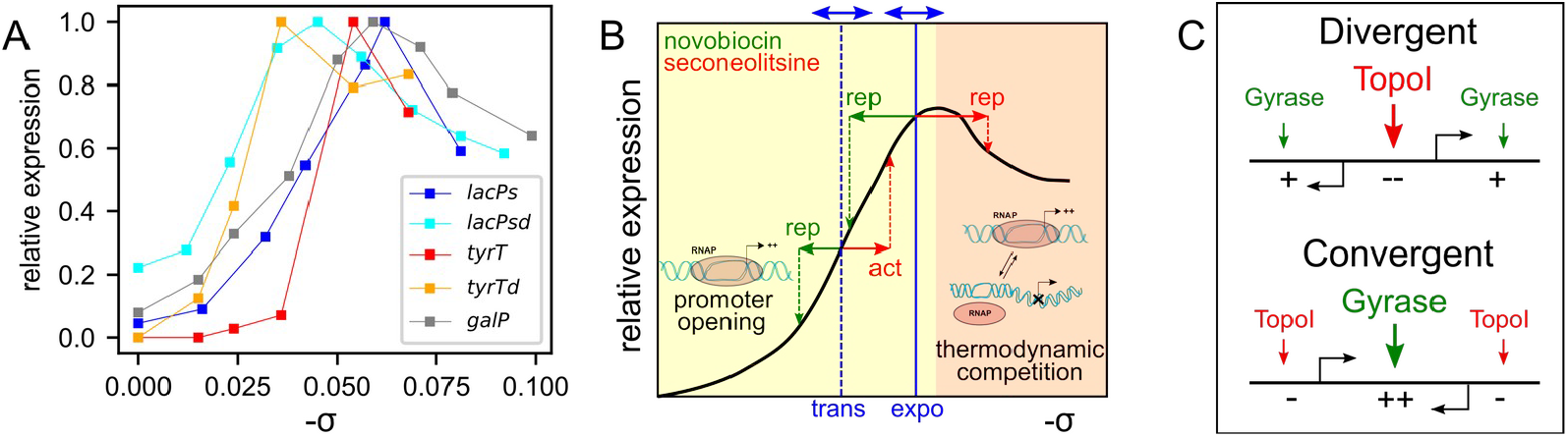
(A) Regulation of various bacterial promoters by SC *in vitro*. The employed native promoters encode either stable (*tyrT*) or messenger RNAs (*galP*), whereas *lacPs* is a mutant promoter derived from *lacP. tyrTd* and *lacPsd* are mutant versions of *tyrT* and *lacPs*, respectively (7, 11, 59). (B) Qualitative regulatory model summarising the data of A (black solid line) with putative mechanisms: promoter DNA opening for open-complex formation (yellow background) (17) and thermodynamic opening competition (orange background, see text) (60). Physiological SC levels valid for many bacteria in exponential or transition to stationary phase are indicated in blue, with double arrows symbolising limited precision and species-dependent variability (61). The expected regulatory effect of an antibiotic shock in either phase is indicated in green. (C) Model of orientation-dependent binding of topoisomerases, and subsequent transcriptional regulation by topoI inhibition (adapted from (30)).

During a relaxation shock, in either phase, the expression rate is predicted to be shifted leftwards to a lower level. The effect of the shock is thus relatively similar in both phases, explaining the comparable pattern of expression observed with novobiocin (Fig. 8B). During a seconeolitsine shock, the situation is different. At the moderate SC level experienced during transition to stationary phase, the SC increase (rightwards shift) induced by topoI inhibition is expected to induce the expression rate of most promoters; in contrast, in exponential phase, the SC level is already close to the maximum of the curve, and the shock thus tends to reduce the expression level of many individual promoters, although this response is likely more sensitive to the exact optimal level of each promoter and thus more complex.

Although very simplified, this analysis from *in vitro* data might thus contribute to several notable observations that we have made from our data: (i) many promoters, such as *lpxC, tonB* and *dps*, respond to novobiocin and seconeolitsine in the same direction (Fig. 11), suggesting a non-monotonous SC-activation curve; (ii) the expression pattern associated to seconeolitsine is more condition-dependent than that of novobiocin (Fig. 8); (iii) seconeolitsine mostly represses promoters in exponential phase, and activates them at the transition to stationary phase (Fig. 6C).

### Role of topoisomerase I in resolving transcription-induced supercoils

While the genome-averaged SC level has a global regulatory effect, several observations made above also further support a specific role of topoI in the handling of negative supercoils generated locally in the wake of elongating RNAPs. The relationship between expression strength and repressive effect of seconeolitsine (Fig. 9) shows that, in the exponential phase, the negative supercoils in that phase (Fig. 12B), and although additional mechanisms (weaker global expression level, nucleoid-associated proteins) might contribute to this effect. The asymmetrical generation of supercoils also results in a relation between the orientation of neighbouring genes and the local induced by the expression of a single gene are sufficient to repress its own expression when topoI is inhibited, and this effect increases with the promoter strength. In contrast, at the transition to stationary phase, strong promoters are activated or repressed in similar proportions, as expected from the more relaxed average level recruitment of topoI (Fig. 12C), which was previously observed by ChIP-Seq (50, 55, 62), and gives rise to the response observed on Fig. 10.

Based on these effects and probably others (in particular related to replication), the picture of a *global* chromosomal SC increase or relaxation induced by the inhibition of either topoisomerase enzyme appears as somewhat misleading, since this inhibition likely rather induces a very heterogeneous modification of *local* supercoils, with a strong effect of the genomic context. This distribution, and its regulatory effects, are likely further complicated by the dynamical repartition of local SC into contributions that affect transcription in different manners, in particular into constrained and unconstrained SC fractions, and into twist and writhe. These contributions are affected very differently by the two main considered topoisomerase enzymes, since DNA gyrase introduces supercoils by crossing two distal loci coming into close spatial proximity, i.e., predominantly introduces writhe (33), whereas topoI cleaves a single strand of negatively supercoiled DNA, i.e., predominantly removes an excess of negative twist (1). They are also strongly affected by the recruitment of nucleoid-associated proteins, most of which induce distortions into DNA and displace the equilibrium between twist and writhe in favour of the latter.

Altogether, a better understanding of this regulation will thus significantly benefit from a detailed and high-resolution mapping of the distribution of local SC levels along the chromosome (63).

## Acknowledgments

We thank Georgi Muskhelishvili for his suggestions and critical reading of the manuscript, Olivier Espéli for useful advice, and Florelle Deboudard for technical help.

