## Supplementary Fig for "What is a supercoiling-sensitive gene? Insights from topoisomerase I inhibition in the Gram-negative bacterium *Dickeya dadantii*"

### Supplementary Information

Maïwenn Pineau, Shiny Martis B., Raphaël Forquet, Jessica Baude, Lucie Grand, Florence Popowycz, Laurent Soulère, Florence Hommais, William Nasser, Sylvie Reverchon and Sam Meyer

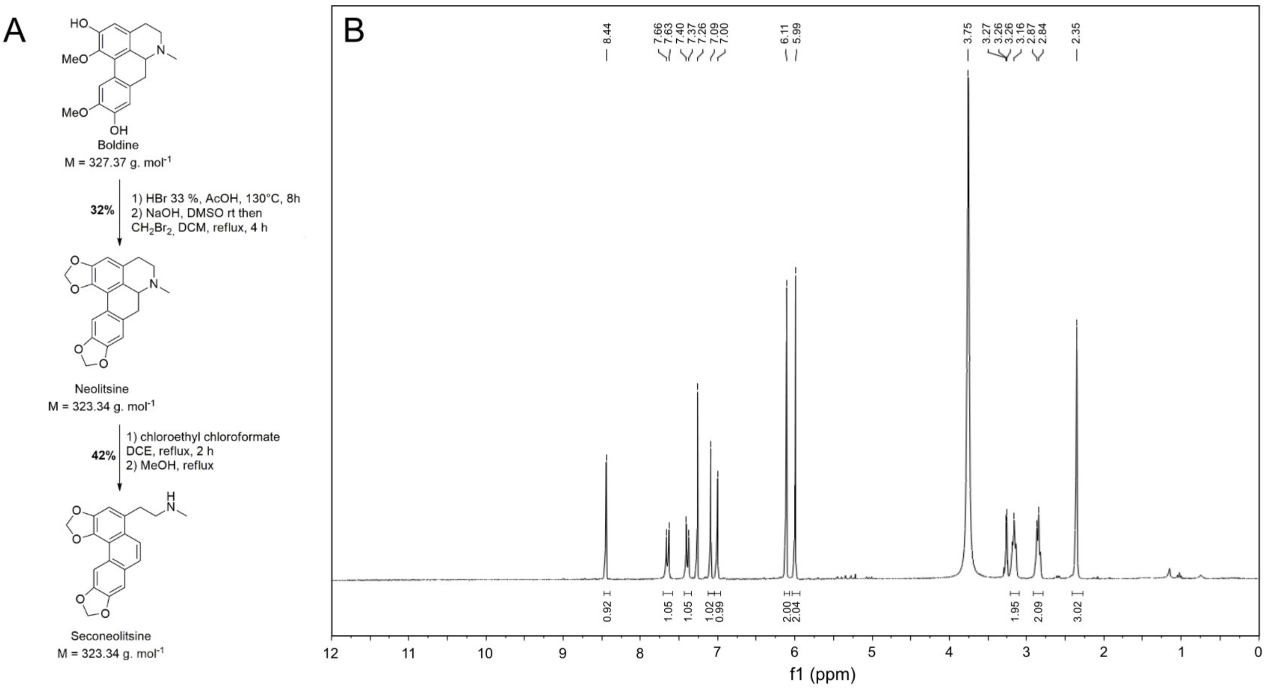


**Fig. S1:** (A) Synthesis of seconeolitsine from boldine. (B) Purity of the product measurement: spectrum of the compound by $^1$H NMR (300MHz in $CDCl_3$ / $CD_3OD$).

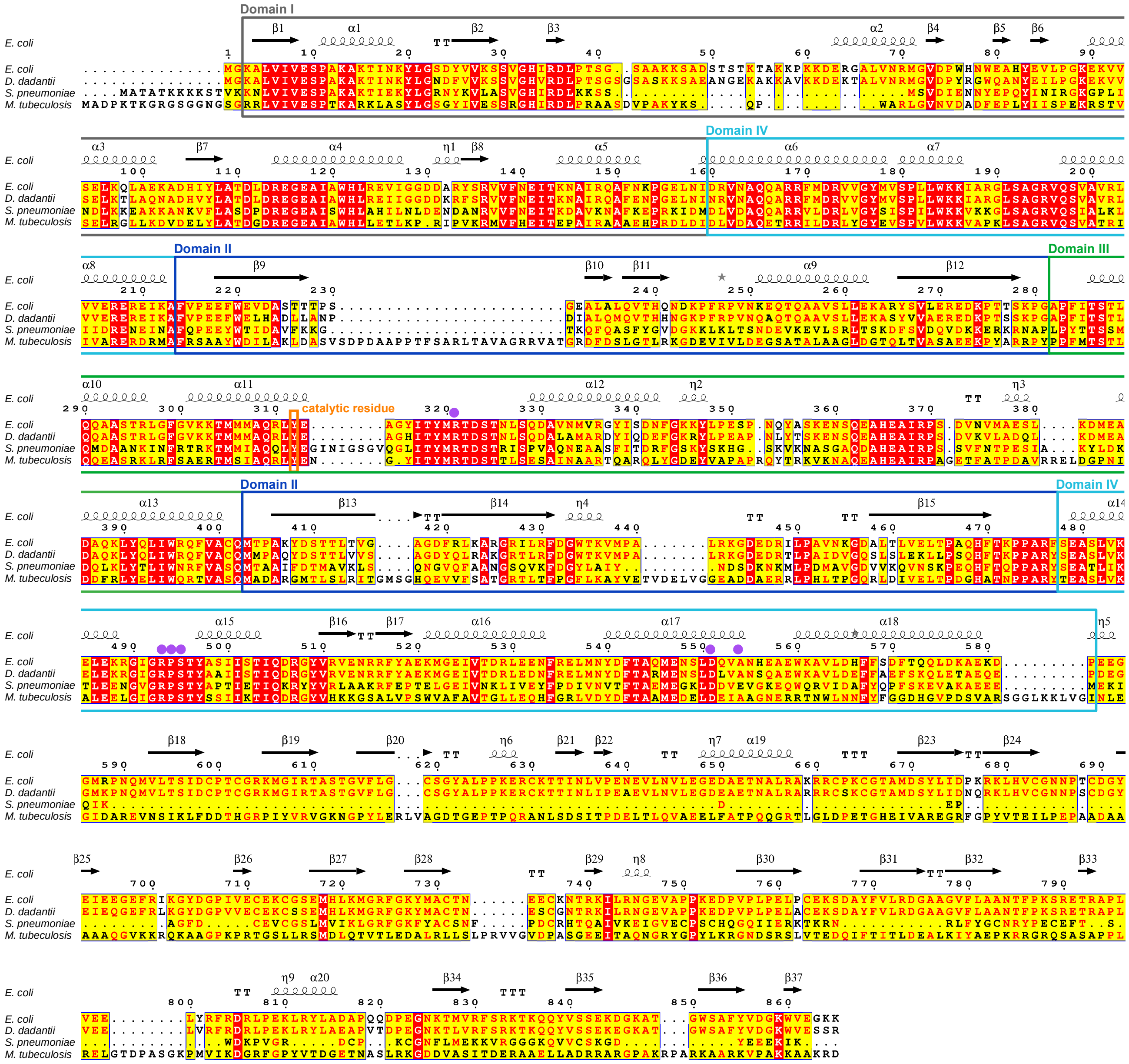

**Fig. S2:** Alignment of *E. coli*, *D. dadantii*, *S. pneumoniae* and *M. tuberculosis* topoisomerase I sequences obtained using ENDscript (1). Identities are highlighted in red, similarities in yellow. Secondary structure elements of *E. coli* topoisomerase I are shown above the sequences (α-helices by squiggles, β-strands by arrows, strict α-turns by TTT letters and β-turns by TT letters). The catalytic tyrosine residue is shown by an orange box (Tyr-319 for *E. coli*). Residues indicated with purple dots are part of the nucleotide-binding site and might interact with seconeolitsine. Grey, dark blue, green and light blue boxes indicate previously described topoisomerase I domains (2).

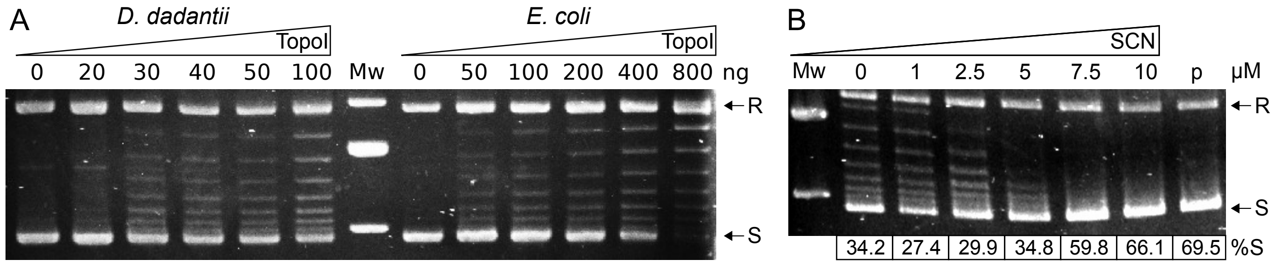


**Fig. S3:** (A) Activities of purified topoisomerase I from *D. dadantii* and *E. coli*. 0.5 µg of pUC18 plasmid was incubated with the indicated mass of topoisomerase I for 15 min at 37°C. R corresponds to pUC18 relaxed form, S to the highly negatively supercoiled form. (B) Inhibition of *E. coli* topoisomerase I by seconeolitsine. The indicated amount of seconeolitsine was first incubated 10 min with 100 ng *E. coli* topoisomerase I at 4°C. 0.5 µg of pUC18 plasmid (p) was then added and the mix was incubated for 15 min at 37°C. %S is the percentage of topoisomers in the highly negatively supercoiled form (S). R indicates the relaxed form.

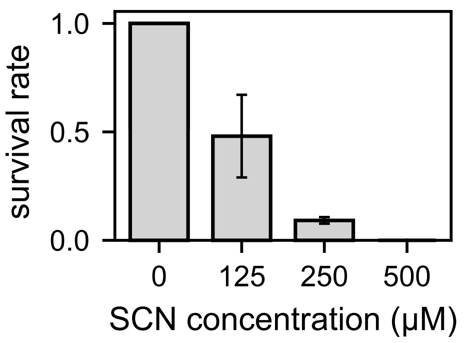


**Fig. S4:** Inhibition of the growth of *E. coli* NM522 cells by seconeolitsine. Each bar indicates the proportion of growing colonies on plates with the specified amount of seconeolitsine compared to plates without seconeolitsine, with a 95% statistical confidence interval.

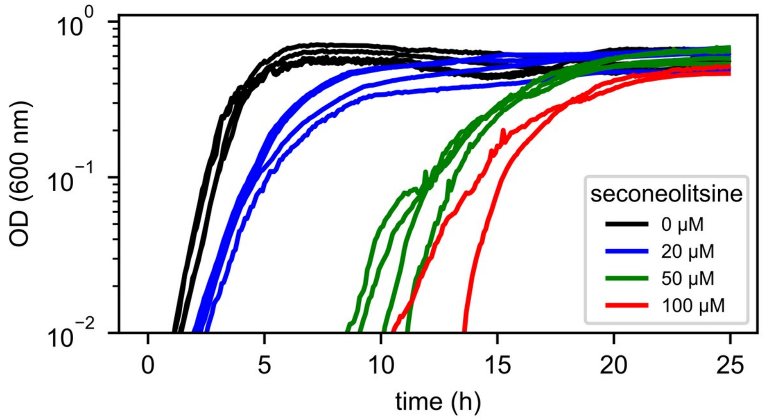


**Fig. S5:** Growth curves of the Gram-positive bacterium *B. subtilis* in presence of increasing amounts of seconeolitsine, solvated in a constant (5% vol) volume of DMSO.

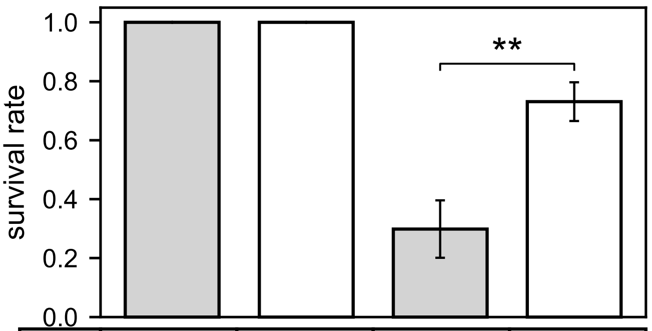


| | | | | |
|---|---|---|---|---|
| SCN | - | - | + | + |
| IPTG | - | + | - | + |
| topoI | + | ++ | + | ++ |

**Fig. S6:** Effect of topoisomerase I expression level on survival rate in presence of seconeolitsine. The *D. dadantii* topoisomerase I gene under the control of an IPTG-inducible promoter was inserted into pQE80L and transformed into *E. coli* NM522 cells. The survival rate is defined as the proportion of growing colonies observed on plates containing 125 µM seconeolitsine vs without seconeolitsine. In the absence of IPTG, *D. dadantii* topoisomerase I is only expressed at a basal level and the survival rate on plates containing seconeolitsine is around 30%. In the presence of 0.5 mM IPTG, *D. dadantii* topoisomerase I is overexpressed and the survival rate is significantly higher (73%, *P*=0.0015).

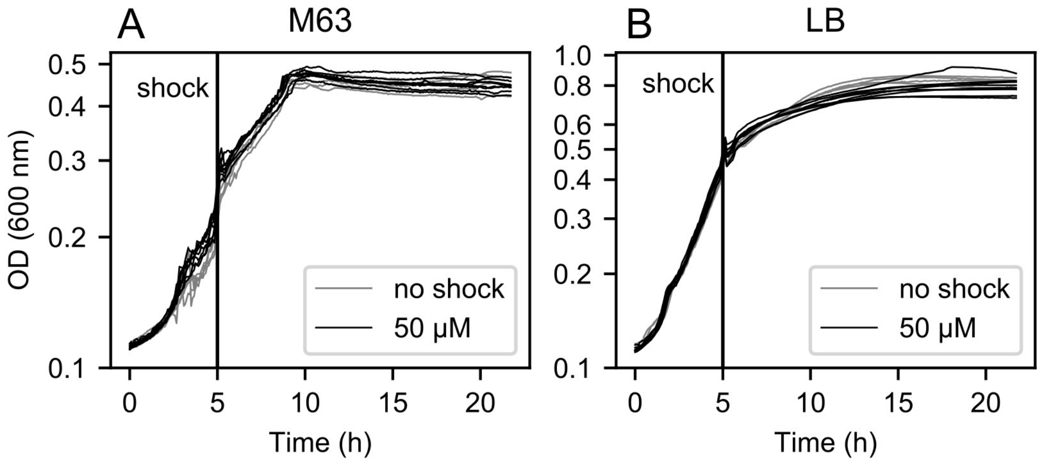


**Fig. S7:** Growth curves of *D. dadantii* treated with a seconeolitsine shock at 50 µM during exponential growth (vertical line), exhibiting no significant effect (multiple black and grey lines indicate replicates of treated and untreated samples, respectively, and are almost indistinguishable). The chosen concentration thus minimises the metabolic effect of the shock, while inducing a significant increase in SC (Fig. 3). The experiment was carried in M63+glucose (A) and LB (B) culture media.

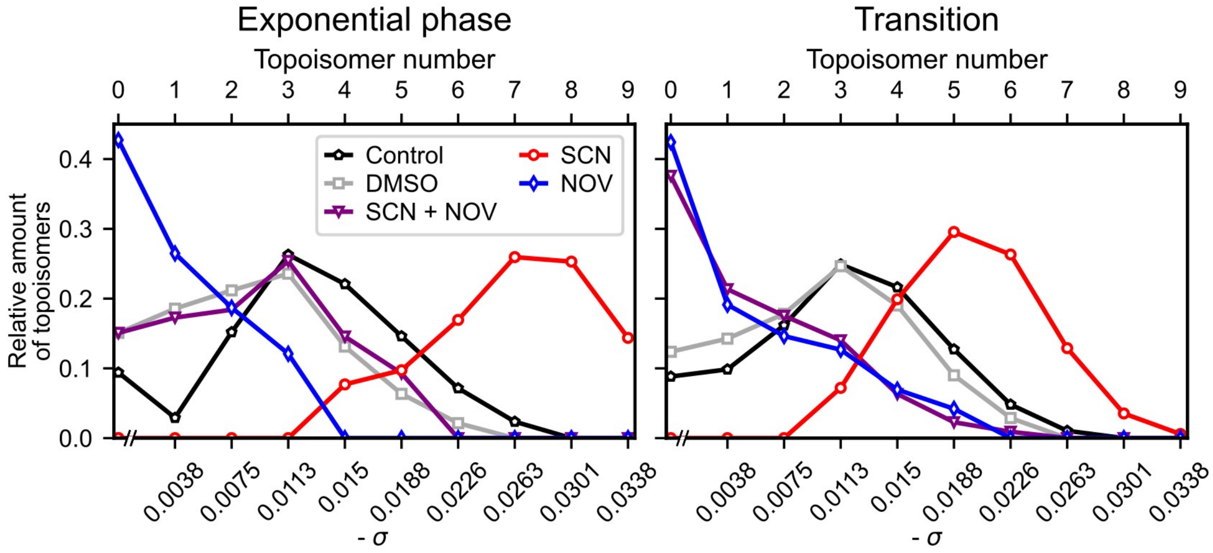


**Fig. S8:** Quantification of topoisomer distribution of pUC18 plasmids isolated from *D. dadantii.* Cells were treated with seconeolitsine (50 µM), novobiocin (100 µg.ml$^{-1}$) or both, in exponential (left) and transition to stationary phase (right). The most relaxed fractions were not fully resolved on the chloroquine gel, hence the discontinuity indicated in the x-axis.

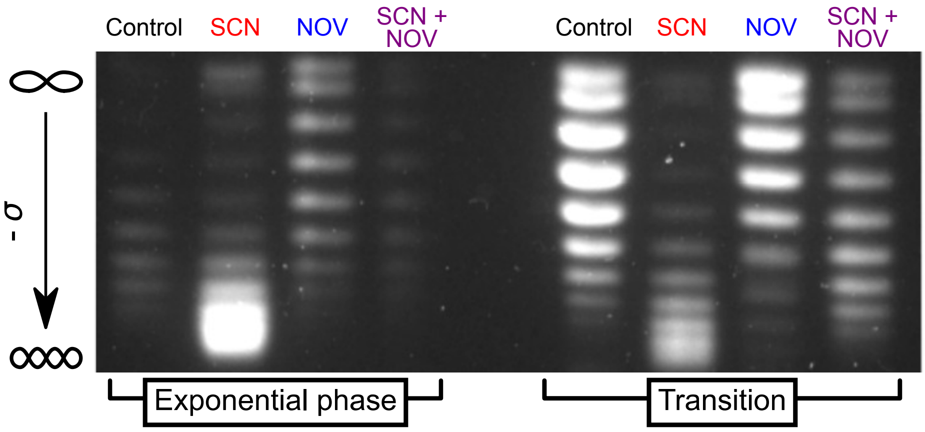


**Fig. S9:** Agarose-chloroquine gels of pUC18 plasmids isolated from *E. coli* NM522 cells in the exponential phase (left) or at the transition to stationary phase (right), 15 min after antibiotic shocks (seconolitsine 50 µM, novobiocin 100 µg.ml$^{-1}$, combination of both, from left to right). The downward migration increases with a negative SC level. The results are qualitatively similar to those obtained in *D. dadantii*, with a stronger effect of seconeolitsine.

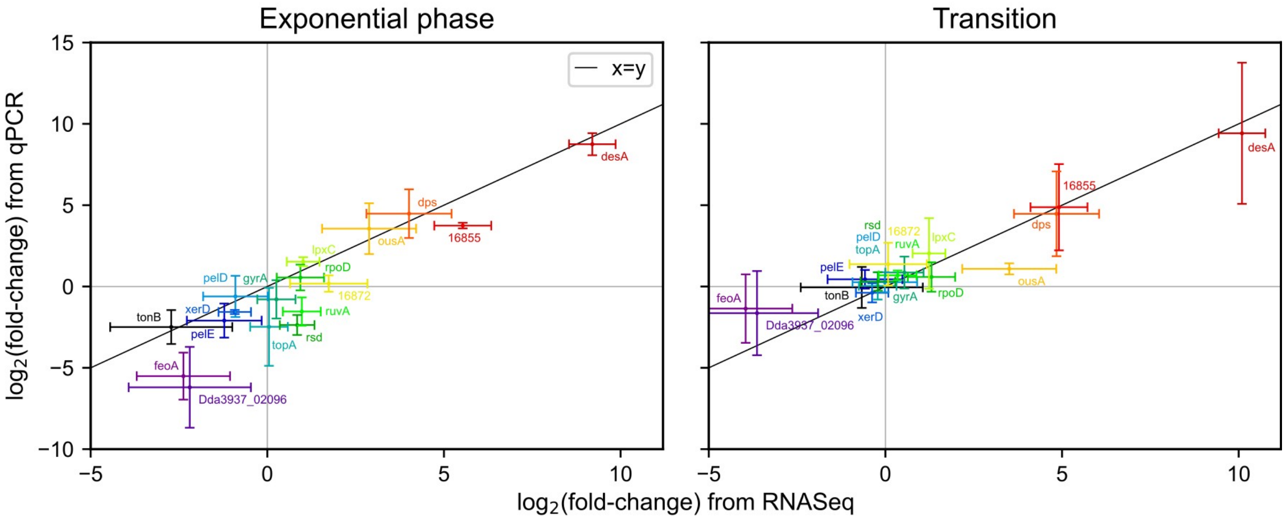


**Fig. S10:** Comparison of gene responses obtained with qRT-PCR and RNA-Seq to seconeolitsine treatment in exponential (left) and transition to stationary phase (right). Pearson's correlation coefficients reached 0.904 with p-value of $6.23e^{-7}$ (exponential phase) and 0.917 with p-value of $2.32e^{-7}$ (transition).

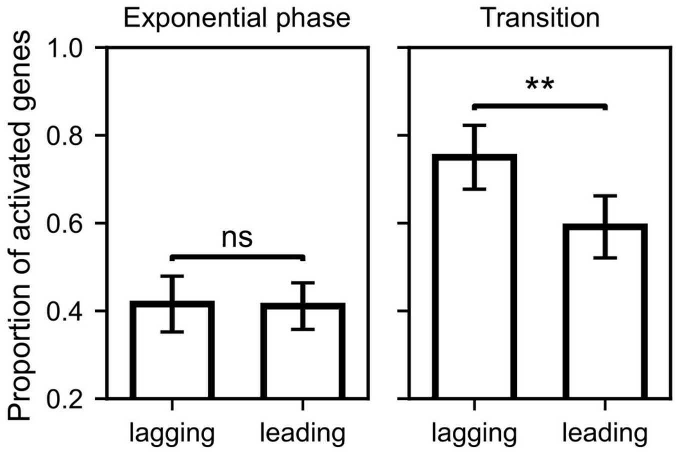


**Fig. S11:** Proportion of activated genes (among differentially expressed ones) on the lagging vs leading replicative strand in exponential phase and at the transition to stationary phase ($P$=0.0015).

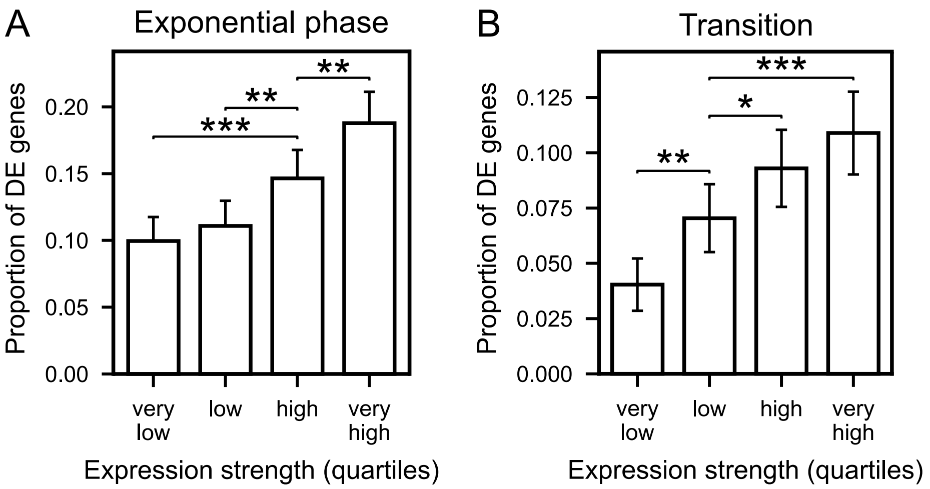


**Fig. S12:** Proportion of differentially expressed genes depending on the expression strength, in the exponential phase (A) or transition to stationary phase (B). Genes were separated into quartiles based on their average number of reads in all samples.

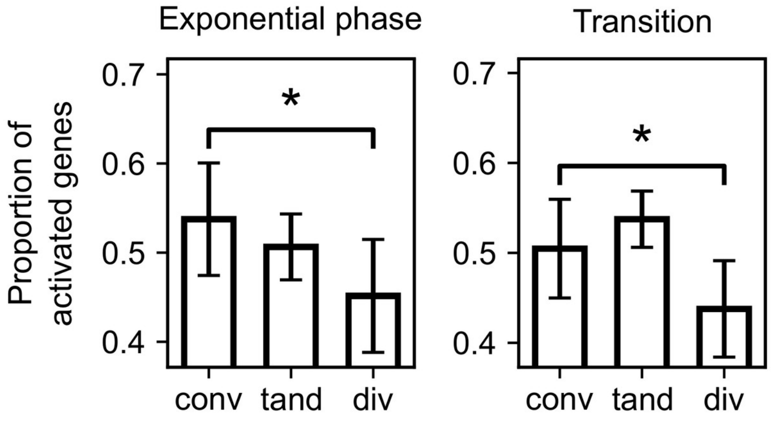


**Fig. S13:** Proportion of activated genes (among differentially expressed ones) on convergent, tandem and divergent genes following novobiocin treatment in *D. dadantii* in exponential phase ($P$=0.030) and at the transition to stationary phase ($P$=0.044) (3).

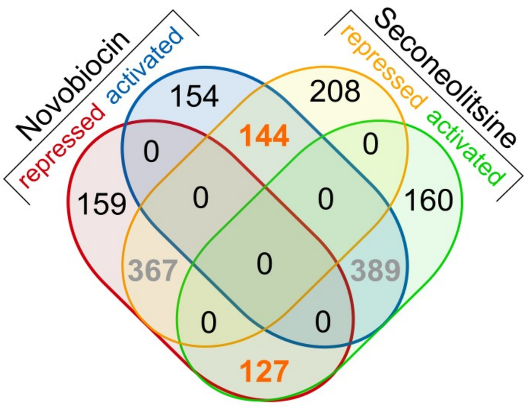


**Fig. S14:** Venn diagram of the response of *S. pneumoniae* genes to novobiocin and seconeolitsine treatments in exponential phase, computed from the data of (4). The number of genes responding in opposite directions to the two drugs are indicated in orange, and those in the same direction in grey.

**Tab. S1:** List of primers used in PCR.

| Gene / plasmid name | Accession number | Forward primer | Reverse primer |
|---|---|---|---|
| *ryhB* | Dda3937_04678 | GACTGAGAATTCGCGTTCCGGGAAAGC CAAA | CATAGCATGCTAAAAAAAAGCCAGCAC CCGAGC |
| *tonB* | Dda3937_00565 | GACGCCAGAACCGGAAATAC | CGTTTCCTTCGGTGGTTCC |
| *ousA* | Dda3937_00791 | GCAACACGTGGAGAAACTGG | CTTGGCATGTAGGTCAGCAG |
| *ruvA* | Dda3937_01621 | CGGCGTCGGATATGAAGTTC | CACGCCGTTCACTTTGATCA |
| *rsd* | Dda3937_00236 | ACGCTGGATGAATTTTGCCA | ATTTGCTGGGTGTTGCTACG |
| *dps* | Dda3937_02884 | ACACGAGATGCTGGATACCT | CGTTGTGGATATCAAGCGGG |
| *xerD* | Dda3937_02296 | GCTGCCGAAAGATCTGACG | TGTTGCCCTTCCCTATCACG |
| *desA* | Dda3937_00786 | CCATCAGCCAGCAAACCTAC | GCGAAGGTCAGATCCAGTTG |
| | Dda3937_01473 | GGCGGAACAGAACCTTAACC | CGCTTTCACGTACGGATACC |
| | Dda3937_01484 | AACGGAACAAGATCAGCTGC | AGGTCATACAGCTCGTTCCA |
| *lpxC* | Dda3937_02232 | ACGGTAAGCGCATTCTGGTA | CATCCAGCAGCAGGTAGACA |
| *pelD* | Dda3937_03372 | TTGTGGAAGGTAACGCGCAGTTTG | ATGGCAAATTCACCAACGGCTCTC |
| *pelE* | Dda3937_03371 | AGCGAATTCAAAGCAGCACT | GGCGTTTCGATGTACAGGTT |
| *topA* | Dda3937_00587 | GAGCAAGAAAAGCGCTGAAG | TTGCGAGATAGACGTGATCG |
| *rpoD* | Dda3937_03534 | CCGACCGCTATTGATGAAAT | CACAGTTCGACGCACAGTTT |
| *gyrA* | Dda3937_01774 | GGTAAATATCACCCGCATGG | TCTCCAGATCGGAAAGCAGT |
| *feoA* | Dda3937_02095 | GGGGTT TCAGAAAGGATCGG | AGAATCAAAATCTGCGCTTCG |
| | Dda3937_02096 | TCAAACGCTGCT TGATCTCC | TTGCCCATCTCCCAGTTG AT |
| *rpoA* | Dda3937_01515 | AAACCGCGCCTGGTAGATA | CCTTTCAGGTTGAGCAGGAT |
| *topA* | Dda3937_00587 | CACCATCACCATCACGGATCCATGGGTA AAGCTCTCGTTATCGTC | AAGCTCAGCTAATTAAGCTTCTCGAGTT AGCGGCTGCTTTCCACCCAC |
| pQE80L | | AAGCTTAATTAGCTGAGCTTGGACTCC | GGATCCGTGATGGTGATGGTGATGCGAT CC |

**Tab. S2:** Lists of *D. dadantii* genes responding to seconeolitsine treatment in exponential and transition to stationary phase.

**Tab. S3:** Lists of *D. dadantii* genes (locus tags and gene names or ASAP identifiers (5)) significantly affected by seconeolitsine and novobiocin treatment.
